# StrucTrace: Fourier Watermarking for Traceable Bio-molecular Assets

**DOI:** 10.1101/2025.10.18.683214

**Authors:** Xu Wang, Tin-Yeh Huang, Yiquan Wang, Yafei Yuan

## Abstract

The rise of generative artificial intelligence (GenAI) in protein and nucleic acid design has created unprecedented opportunities for synthetic biology, but also heightened the need for reliable provenance and intellectual-property protection. To meet this challenge, we present a Fourier domain watermarking framework that encodes digital identifiers directly into three-dimensional biomolecular structures. By perturbing only flexible backbone atoms and embedding information through frequency domain modulation, the method achieves imperceptible alterations while ensuring deterministic and reversible decoding. Large-scale validation on over 40,000 protein structures demonstrates its robustness: structural deviations remain orders of magnitude below biological thresholds, watermarks are recovered with perfect accuracy, and functional analyses confirm stability at both thermodynamic and dynamic levels. Beyond technical performance, the approach provides a foundation for a broader ecosystem of secure biomolecular asset management, integrating provenance verification, access control, and digital rights management. Together, these advances establish biomolecules as traceable and auditable digital assets, aligning the future of bio-design with emerging standards for trustworthy AI.

## 1 Introduction

Generative artificial intelligence (GenAI) is accelerating discovery in synthetic biology by enabling the rapid creation of novel proteins and nucleic acids [1–4]. However, it also presents a critical challenge: ensuring traceability. As these molecules become structurally indistinguishable from their natural counterparts, verifying their origin is essential for intellectual-property attribution [5, 6], regulatory compliance [7], and biosecurity [8–10].

While combining GenAI with biomacromolecule watermarking is a promising research direction, current methods face several fundamental limitations. First, their applicability is restricted, as they are often tightly coupled to specific generative models [11–13] and cannot be used on experimentally derived structures. Second, their “black-box” nature hinders interpretability, preventing a transparent assessment of any structural perturbations. These pose a practical challenge to collaboration, as the verification process often requires full molecular disclosure, which conflicts with data privacy requirements [14, 15].

To overcome these challenges, we introduce StrucTrance (**Struc**ture **Trace**), a watermarking framework that directly embeds digital identifiers into 3D biomolecular structures. By applying a Fourier transform to atomic coordinates, the method creates a deterministic geometric encoding, confining perturbations to targeted regions to preserve functional integrity. This approach ensures unambiguous watermark retrieval, achieving 100% bit accuracy while reducing structural deviation 300-fold (scRMSD) compared to state-of-the-art methods. StrucTrance thus establishes a robust foundation for auditable and privacy-preserving biomolecular design, in line with emerging trustworthy AI standards.

## 2 Methods

We introduce a frequency-domain method to embed digital watermarks into 3D protein structures by applying imperceptible coordinate perturbations to structurally flexible C_*α*_ atoms. This process, designed to preserve biological function, is illustrated in Figure 1.

**Figure 1.**
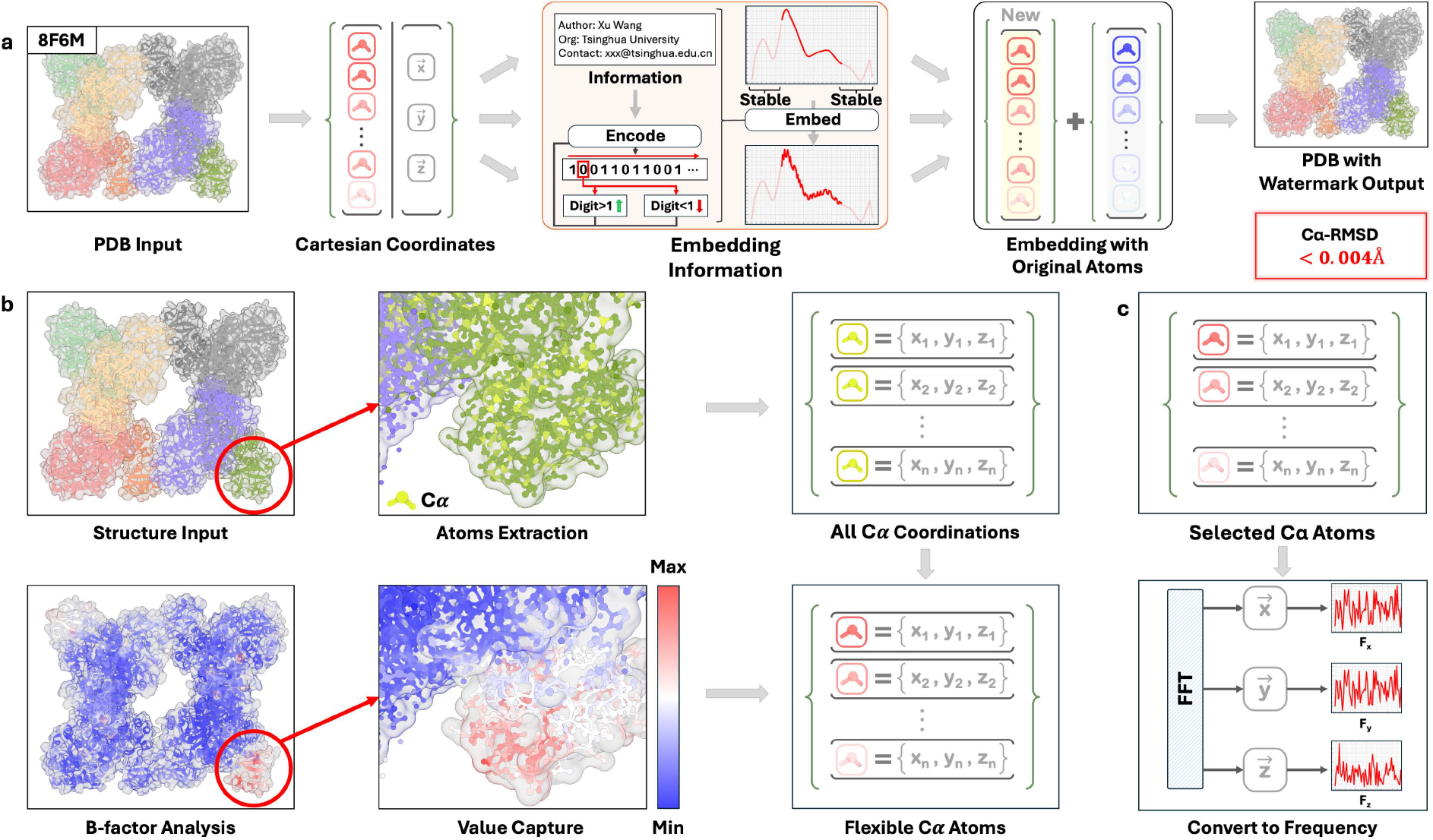
Workflow for frequency-domain digital watermarking of protein structures. **(a) The overall watermarking pipeline.** Starting with an input PDB file, the 3D coordinates of target C_*α*_ atoms are extracted. These coordinates are then transformed into the frequency domain using a discrete Fourier transform (DFT) for watermark embedding. This frequency range is optimal as it preserves the low-frequency signals that define the global fold while offering greater robustness to noise than high-frequency signals. Phase information is preserved and conjugate symmetry is enforced to ensure the inverse transform yields real-valued coordinates. Then, an inverse DFT (IDFT) converts the frequency-domain signal back into real-valued spatial coordinates, generating a new PDB file that contains the watermark while preserving all original metadata. **(b) Target atom selection based on B-factors**. To preserve the protein’s biological function, perturbations are confined to backbone C_*α*_ atoms in structurally flexible regions. These atoms are identified by their high B-factor values in the source PDB file. Their inherent mobility allows for minor coordinate shifts to be tolerated without compromising the protein’s global fold or the conformation of its active sites. **(c) Watermark encoding via Fast Fourier Transform (FFT)**. The coordinate vectors (*x, y, z*) of the selected C_*α*_ atoms are transformed into the frequency domain. A binary watermark is embedded by modulating the amplitudes of specific mid-range frequency components.

The watermarking algorithm proceeds as follows:

### 2.1 Target Atom Selection

We identify C_*α*_ atoms in structurally flexible regions based on their B-factor values from the input PDB file. High B-factors indicate thermal mobility, allowing coordinate perturbations to be tolerated without disrupting the global fold or active site geometry.

### 2.2 Frequency Domain Transformation

The coordinate vectors (*x, y, z*) of selected atoms are independently transformed using the Discrete Fourier Transform (DFT):

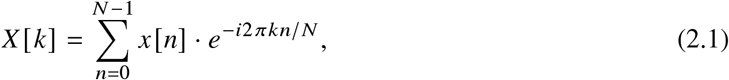

where *N* is the number of selected atoms, *x*[*n*] represents the spatial coordinates, and *X*[*k*] denotes the frequency-domain representation.

### 2.3 Watermark Embedding

A binary watermark *W* = {*w*_1_, *w*_2_, …, *w*_*M*_} is embedded by modulating the amplitudes of mid-range frequency components. Specifically, for each bit *w*_*j*_, we modify the amplitude at frequency index *k*_*j*_ :

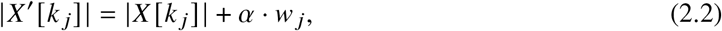

where *α* is a scaling factor controlling embedding strength. Phase information is preserved to ensure the inverse transform yields real-valued coordinates. Conjugate symmetry is enforced to maintain the real-valued property of spatial coordinates.

### 2.4 Inverse Transformation

The modified frequency-domain signal is converted back to spatial coordinates via the Inverse DFT (IDFT):

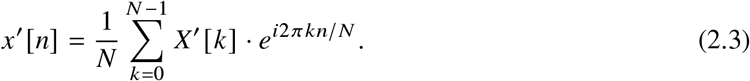

The resulting coordinates *x*′ [*n*] form the watermarked structure, which is exported as a new PDB file with all original metadata preserved.

### 2.5 Watermark Decoding

The decoding process is deterministic and does not require access to the original structure. We re-select the same C_*α*_ atoms using B-factor criteria, apply DFT, and extract the watermark by analyzing amplitude patterns at the targeted frequency indices. Bit recovery is achieved by thresholding amplitude differences against expected embedding patterns.

## 3 Results

We validated our watermarking method across over 40,000 RCSB PDB structures, achieving three key breakthroughs (Tab. 1).

**Table 1.**
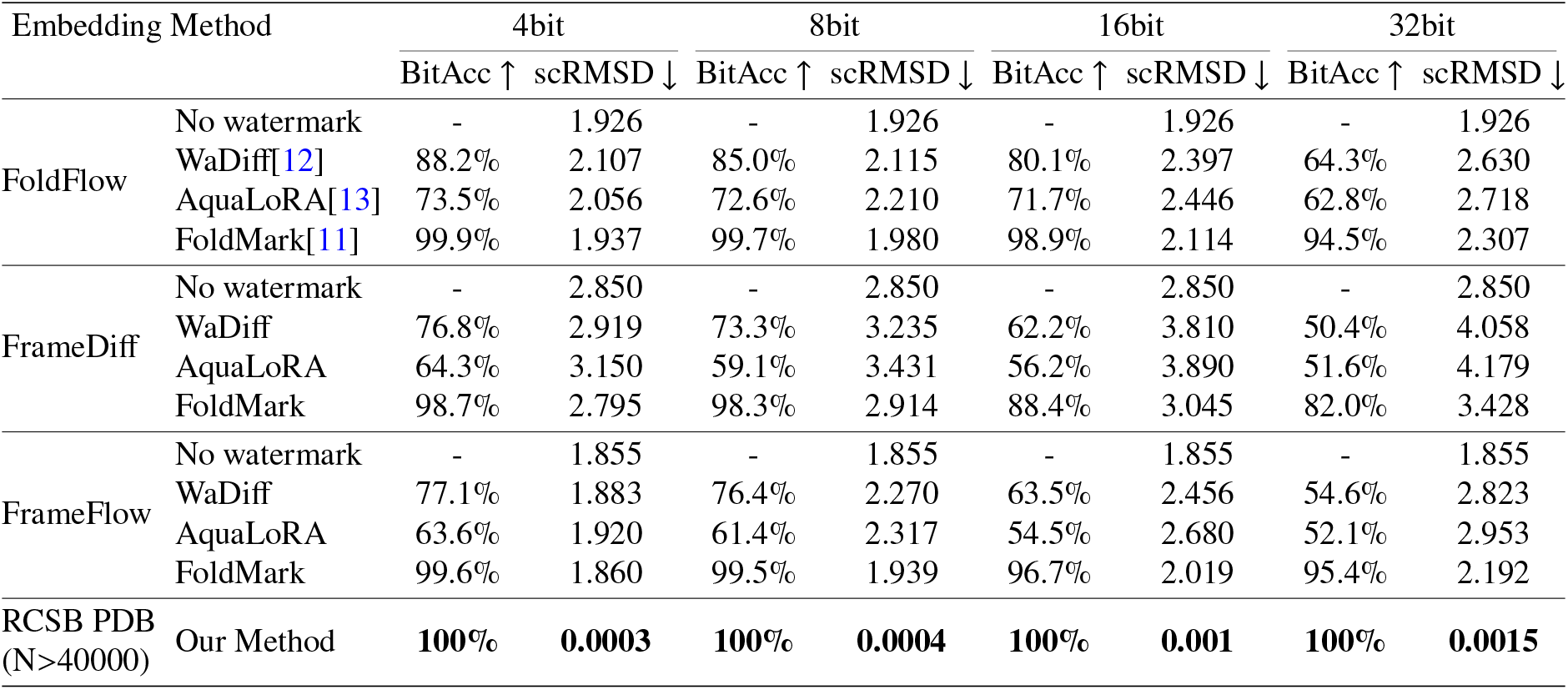
Cross-Method Comparison of Bit Accuracy and Structural Deviation.

First, ultra-low structural perturbation was demonstrated with scRMSD ≤ 0.0015 Å, significantly lower than the structural biology threshold of 2.0 Å for indistinguishable folds. This ensures watermarked proteins remain visually and computationally identical to their originals, as confirmed by alignment tools and manual inspection.

Second, high-capacity information embedding was achieved with 100% bit accuracy (BitAcc) for 4– 32 bit payloads. Structural deviation (scRMSD) increased by only 0.0003 to 0.0015 Å across payload scales, a 300-fold improvement in stability compared to existing AI methods [11–13]. This decouples information density from structural distortion, overcoming a critical limitation of prior approaches.

Third, thermodynamic and dynamic analyses confirmed functional preservation. As Rosetta’s ddg_monomer protocol showed ΔΔ*G* ≈ 0 across all cases, there was no indication of destabilization. Molecular dynamics simulations (50–100 ns) revealed stable backbone RMSD trajectories without unfolding or large-scale drift, with RMSF analysis localizing perturbations to non-functional loop regions.

The method’s universal applicability was tested on diverse systems, from monomeric actin to multi-subunit RNA polymerase II complexes, with consistent performance. It outperformed state-of-the-art models while requiring no database-specific training.

This combination of minimal structural noise, scalable information density, and preserved biological function establishes a new standard for imperceptible watermarking in bio-molecular assets, addressing IP protection needs in structural and synthetic biology.

## 4 Discussion

This work introduces a significant advancement in biomolecular intellectual property protection by embedding information directly into the molecular framework. Our approach achieves functional imperceptibility, creating an intrinsic link between the watermark and the molecule without compromising biological function. To translate our watermarking technology into practice, we propose a three-tiered ecosystem for comprehensive biomolecular asset management. This framework, detailed in Figure 2, is designed to address the distinct needs of academic provenance, industrial security, and commercial licensing for bio-designs.

**Figure 2.**
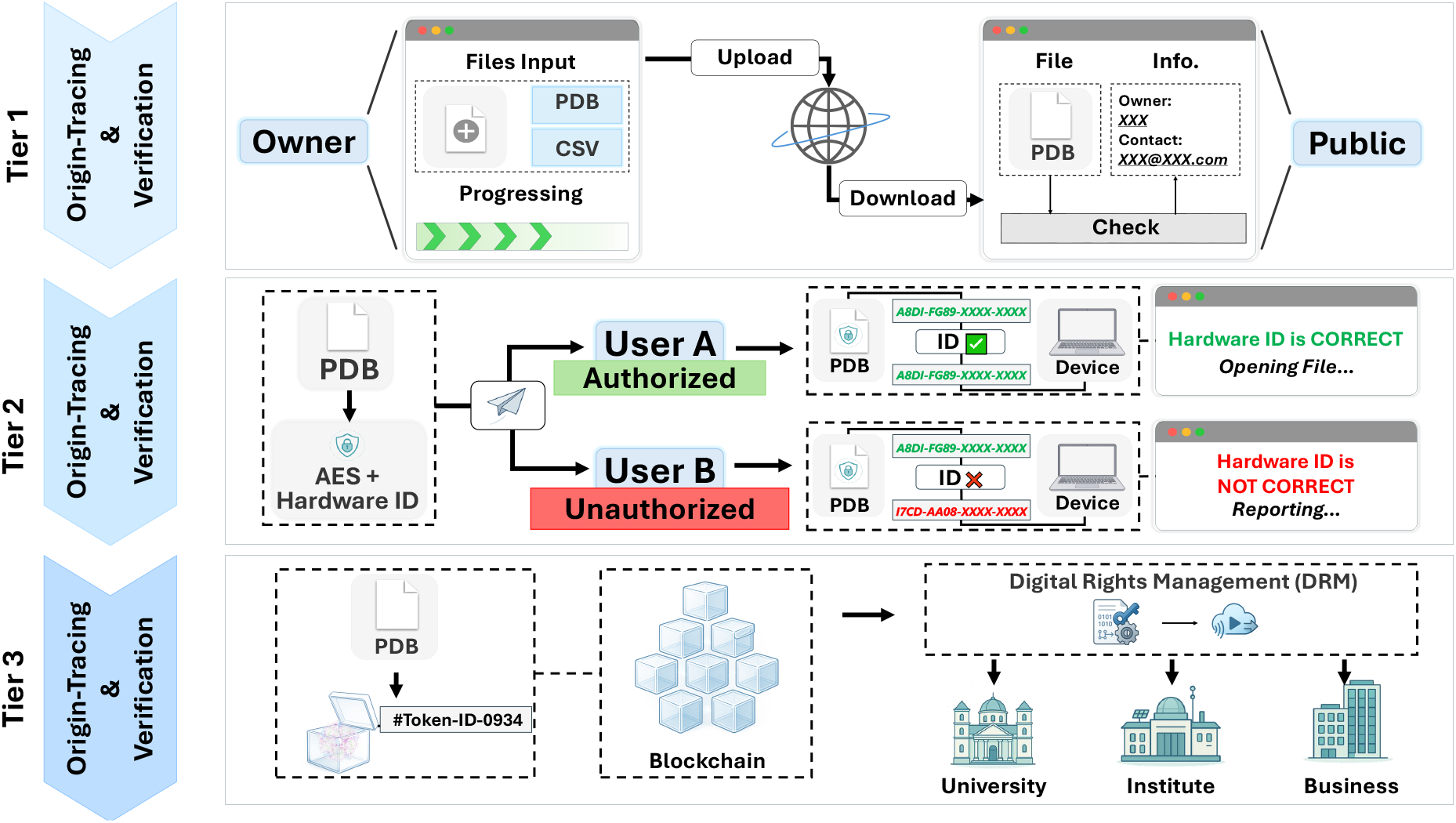
A three-tiered framework for biomolecular provenance, access control, and digital rights management. Our watermarking technology underpins a tiered system for managing biomolecular assets across different use cases. **(Tier 1) Provenance Verification for Public Release**. The watermark links a structure to verifiable metadata, establishing a public record of intellectual priority for academic and open-source applications. **(Tier 2) Hardware-Bound Secure Access**. For proprietary industrial designs, the watermark functions with an encryption layer to restrict access to authorized hardware. Unsuccessful access attempts are logged, providing a security audit trail. **(Tier 3) Blockchain-based Digital Rights Management (DRM)**. To facilitate a commercial market, each watermarked structure is assigned a unique blockchain token. This token represents ownership and enables transparent, licensed distribution and rights management on a decentralized ledger.

Ultimately, this integrated system provides the cornerstone of trust for the burgeoning bioeconomy. As AI-driven protein design proliferates, it addresses the urgent need for provenance in both academic research and industrial collaborations. By establishing the technical foundation for structure-embedded intellectual property, our work redefines biomolecules as secure, verifiable, and tradable digital assets. This framework fosters innovation by ensuring proper attribution for creators, paving the way for a more transparent and equitable future in biotechnology. The system has been deployed at our center, securing our entire database and our flagship peptide design competition.

## Acknowledgments

We thank the Beijing Frontier Research Center for Biological Structure, Tsinghua University, for their support and assistance with this research.

